# Food interactions observed in a pharmacokinetic investigation comparing two marketed cold preparations (BNO1016 and ELOM-080) after administration to male beagle dogs

**DOI:** 10.1101/2021.08.13.456103

**Authors:** Jan Seibel, Astrid Neumann, Anne Müller, Meinolf Wonnemann

## Abstract

The cold remedies Sinupret extract (BNO 1016) and Gelomyrtol forte (ELOM-080) represent the two top-selling cold remedies in Germany nowadays. Whereas BNO 1016 is a typical immediate release coated tablet, ELOM-080 is an enteric-coated soft gelatin capsule. The latter formulation, however, is at risk of pharmacokinetic interactions affecting absorption especially in the case of concomitant food intake. In the present pilot study, we investigated the risk of a possible food effect on BNO 1016 in comparison to ELOM-080 in three male beagle dogs. Single doses of BNO 1016 and ELOM-80 at 80 mg/kg and 160 mg/kg, respectively, were administered under fasting and fed conditions according to a 4-period, 4-treatment within-design. Blood sampling took place until up to 30 h *p.a.* and plasma concentrations of gentiopicroside, verbenalin and from a further marker analyte (BNO 1016 analytes) as well as 1,8-cineole, limonene and perillic acid (ELOM-080 analytes) were determined. Pharmacokinetic parameters focusing on rate and extent of absorption were derived. BNO 1016 analytes demonstrated a homogenous course in all animals in both, the fasted and fed state. Pharmacokinetic characteristics of a typical immediate release drug formulation were observed for all analytes and a food effect could be ruled out. ELOM-080 analytes also showed a homogeneous picture in the fasted state. However, lag-times (t_lag_) of up to 2 h *p.a.* with corresponding t_max_ values of 3 to 4 h were observed, reflecting a longer gastric residence time of the formulation. In the fed state, ELOM-080 showed significant pharmacokinetic characteristics suggesting a clear food effect. A major observation was a double peak phenomenon that could be observed in two out of three dogs. Furthermore, lag-times of some analytes up to 3-4 h and corresponding t_max_ values of up to 6-8 h occurred. In contrast to BNO 1016, these findings suggest that, as with other enteric-coated formulations, there may be a significant risk for food effects with ELOM-080 also in humans.

## Introduction

Worldwide, over-the-counter herbal remedies are clearly dominated by oral dosage forms. The spectrum ranges from immediate-release tablets, capsules, lozenges to effervescent tablets, solutions, syrups, and teas. It is noticeable that modified-release or sustained-release dosage forms obviously do not play a significant role in phytotherapy, although therapeutically they could provide a good benefit especially in some chronic indications [1][2]. Looking at herbal cold preparations, products for the treatment of sinusitis/rhinosinusitis and acute or chronic bronchitis dominated the market in Germany in 2020 [3]. The two top-selling cold remedies (R05C class) in pharmacies were Sinupret^®^ extract (BNO 1016) and Gelomyrtol^®^ forte (ELOM-080), with around 3 million packs sold. They are both marketed also internationally and in member states of the European Union. Surprisingly, ELOM-080 is a modified-release dosage form, more precisely an enteric-coated soft gelatin capsule. In general, the purpose of enteric-coated medicines is either to protect the contained drug from *e.g.* degradation until it is released in the intestine, or to protect the stomach from damage by the drug. One reason for the dosage form of ELOM-080 could be that it is a preparation with essential oil components that perfectly fit to soft gelatin capsules. The special essential oil distillate ELOM-080 [4] has been shown to activate mucociliary clearance [5][6][7]. This has also been proven for BNO 1016, an immediate-release coated tablet formulation [8][9][10], containing an extract of cowslip flowers, yellow gentian root, black elderberry flowers, sorrel herb and verbena herb [11]. Moreover, non-clinical data have demonstrated its anti-inflammatory activity [12]. Flavonoid compounds are being discussed as the active ingredients. Since both preparations are used for acute infections of the respiratory tract, the dosage forms should be designed in such a way that a rapid release of the active substance and a rapid therapeutic effect can be achieved.

Modified release or enteric-coated oral dosage forms, however, are at risk of pharmacokinetic interactions affecting the absorption process especially in the case of concomitant food intake [13][14]. Such food effects are often not predictable from the *in vitro* characteristics of a dosage form. Several investigations have demonstrated a high probability of food interactions for formulations with pH-dependent release properties, such as enteric-coated formulations [16], which present quite sizeable, indigestible solid particles. Passage of such particles through the pylorus occurs normally during phase III of the migrating motor complex (MMC). Food intake interrupts the MMC and can thereby delay the passing of enteric-coated formulations from the stomach into the duodenum [17][18]. Therefore, an *in vivo* food interaction might also be suggested to occur with the intake of ELOM-080 enteric-coated capsules.

### Aim of the study

In the present study we compared possible effects of food intake on ELOM-080 enteric-coated capsules and BNO 1016 immediate release coated tablets in male beagle dogs. Beagle dogs are considered a standard model to predict possible food effects of drug formulations [18].

## Methodology

### Test items and dose selection

For the experiments, commercially available Sinupret^®^ extract coated tablets and Gelomyrtol^®^ forte soft gelatin capsules containing BNO 1016 or ELOM-080, respectively, were purchased in a public pharmacy. The 5-fold equivalent of the recommended human daily dose was chosen (BNO 1016: 5 tablets per animal, *i.e.* 80 mg/kg, ELOM-080: 6 capsules per animal, *i.e.* 180 mg/kg) to ensure bioanalytical detectability of the selected analytes in plasma.

### Study design and animal treatment

The study was approved by the Avogadro LS Animal Ethics Committee. Animal housing and care complied with the recommendations of Directive 2010/63/EU.

Three non-naïve male adult beagle dogs (Marshall BioResources, Lyon, France) weighing 10.9 to 11.4 kg, collectively housed in pens, were used in the study which was performed according to a 4-period, 4-treatment within-design. Animals received all treatments under both fasted (periods 1 and 3) and fed (periods 2 and 4) conditions. They were fasted overnight in each period for at least 8 h prior to dosing. In periods 1 and 3, a high fat diet was given about 3 hours after administration (fasted state). In periods 2 and 4, the same high fat diet was given approximately 10 minutes prior to dosing. Then, regular feeding (Teklad 2027, Envigo Teklad Diets, Madison, USA) was given about 12 hours post drug administration (*p.a.*) in each period. Water was offered *ad libitum*. Treatments were separated for at least 7 treatment free days to guarantee a sufficient washout of the drugs.

Test items were administered orally directly into the back of the throat of the animals. 10 mL of water was added directly into the mouth during administration to ensure good esophageal transit and then 50 mL was administered by gavage using a gastro-esophageal tube to mimic ingestion with liquid in humans as recommended.

### Blood sampling

Blood samples (2.5 mL) were withdrawn into lithium heparin tubes from the jugular vein. Samples were collected at the following time points: pre-dose and 0.5, 1, 2, 3, 4, 6, 8, 12, 18, 24, 26, 28, and 30 h *p.a*.. Real blood sampling times were noted.

### Bioanalytics and pharmacokinetics

After withdrawal, blood samples were immediately centrifuged at 2500 × g for 10 min at +5°C and plasma obtained was cooled in dry ice. Samples were sent on dry ice and stored at ≤ − 70°C until measurement. A maximum period of 8 weeks elapsed from blood withdrawal to measurement. Reliably detectable and representative ingredients or metabolites were selected for measurement: 1,8-cineole, limonene, and its metabolite perillic acid were determined for ELOM-80, gentiopicroside, verbenalin, and for a further marker analyte for BNO 1016. This further analyte is considered a standard marker substance for BNO 1016, but shall not be described in more detail in here for internal company reasons. Qualified and validated methods were used for the analytical measurements. For monoterpene analysis plasma samples were processed by solid phase extraction. 1,8-cineole and limonene were quantitatively deter-mined by GC-MS. The monoterpene metabolite perillic acid as well as gentiopicroside, verbenalin, and the further marker analyte were determined after protein precipitation by LC-MS/MS. Calibration ranges were 20-10000 ng/mL for 1,8-cineole and 20 – 5000 ng/mL for limonene. Perillic acid had a range of 100-9000 ng/mL, gentiopicroside and verbenalin a range of 5-1000 ng/mL and the further marker analyte a range of 2-400 ng/mL. The lower limit of quantitation (LLOQ) is defined as the lowest concentration of each calibration range. Calibration ranges all covered the analytical plasma concentrations of the study samples. Calibration and quality control samples were prepared in canine plasma. QC sample accuracies were found within (gentiopicroside, verbenalin, marker analyte) or slightly outside the range of 100 ± 15% (83.43-95.79% for perillic acid, 84.95 % - 109.3 % for 1,8 cineole, 83.01 % - 143.8 % for limonene.

Pharmacokinetic parameters derived were determined model-independently using Certara Phoenix 64 WinNonlin software (version 8.2.0.4383). Plasma concentrations below LLOQ were set to “0” before C_max_, afterwards omitted. C_max_ and t_max_ values were read directly from the observed concentration-time points. Areas under the curves were calculated according to the linear/log trapezoidal rule, which uses linear data obtained during the absorption phase up to C_max_ and logarithmically transformed concentrations thereafter. The apparent terminal elimination half-life was calculated using non-linear regression on data points visually assessed to be on the terminal phase of concentration curves. Half-life times were only determined when meaningful. The lag-time was taken as the time interval from dosing to the sampling time point for the first quantifiable concentration of an analyte.

## Results

Overlays of individual plasma concentration *vs.* time curves of dogs receiving BNO 1016 under fasting and fed conditions are given in Figure 1 A and B, of dogs receiving ELOM-080 in Figure 2 A and B. The corresponding mean curves are depicted in Figure 3 and 4 and the pharmacokinetic evaluation is given in Table 1.

**Table 1:**
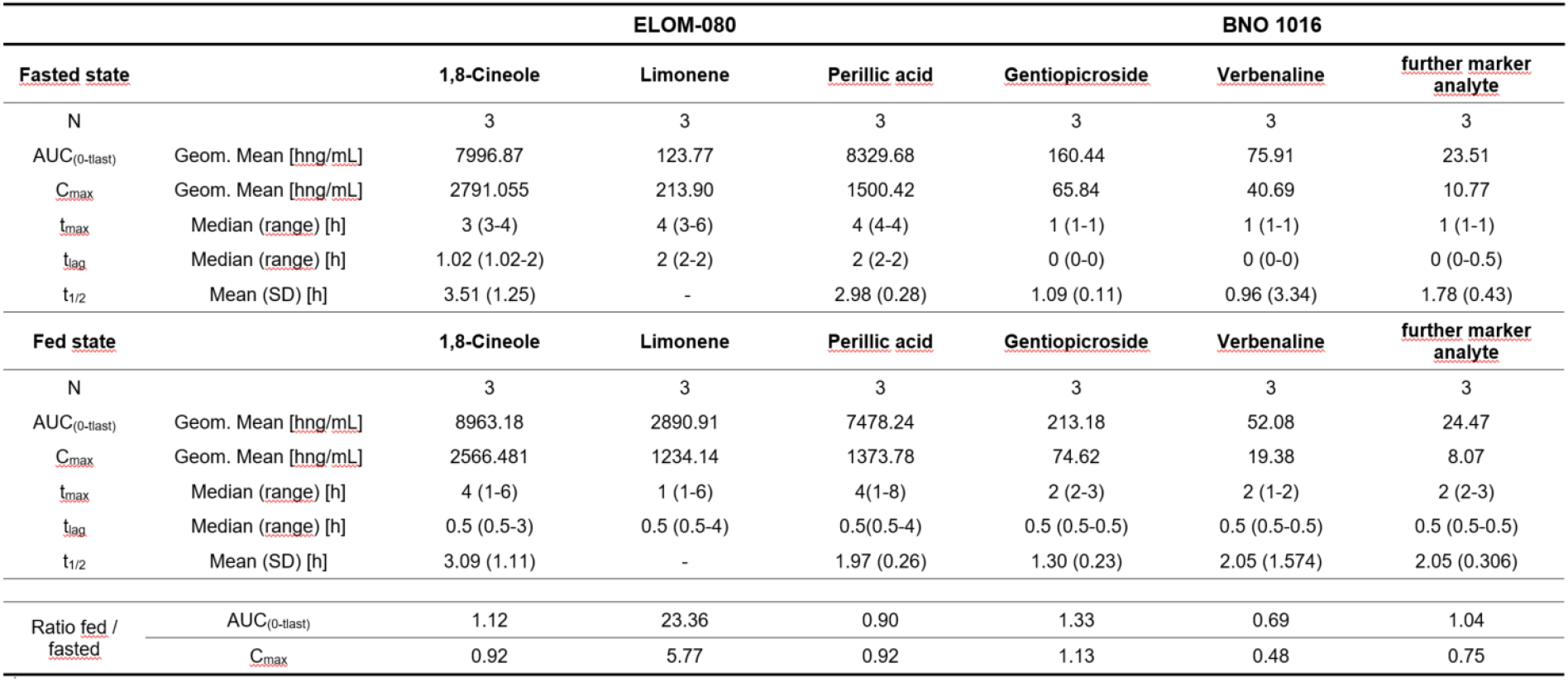
pharmacokinetic parameters after administration of ELOM-080 and BNO 1016 in the fasted and fed state

|  |  | ELOM-080 |  |  | BNO 1016 |  |  |
| --- | --- | --- | --- | --- | --- | --- | --- |
| <u>Fasted state</u> |  | <u>1,8-Cineole</u> | <u>Limonene</u> | <u>Perillic acid</u> | <u>Gentiopicroside</u> | <u>Verbenaline</u> | <u>further marker analyte</u> |
| N |  | 3 | 3 | 3 | 3 | 3 | 3 |
| AUC<sub>(0-last)</sub> | Geom. Mean [hng/mL] | 7996.87 | 123.77 | 8329.68 | 160.44 | 75.91 | 23.51 |
| C<sub>max</sub> | Geom. Mean [hng/mL] | 2791.055 | 213.90 | 1500.42 | 65.84 | 40.69 | 10.77 |
| t<sub>max</sub> | Median (range) [h] | 3 (3-4) | 4 (3-6) | 4 (4-4) | 1 (1-1) | 1 (1-1) | 1 (1-1) |
| t<sub>lag</sub> | Median (range) [h] | 1.02 (1.02-2) | 2 (2-2) | 2 (2-2) | 0 (0-0) | 0 (0-0) | 0 (0-0.5) |
| t<sub>1/2</sub> | Mean (SD) [h] | 3.51 (1.25) | - | 2.98 (0.28) | 1.09 (0.11) | 0.96 (3.34) | 1.78 (0.43) |
| <u>Fed state</u> |  | <u>1,8-Cineole</u> | <u>Limonene</u> | <u>Perillic acid</u> | <u>Gentiopicroside</u> | <u>Verbenaline</u> | <u>further marker analyte</u> |
| N |  | 3 | 3 | 3 | 3 | 3 | 3 |
| AUC<sub>(0-last)</sub> | Geom. Mean [hng/mL] | 8963.18 | 2890.91 | 7478.24 | 213.18 | 52.08 | 24.47 |
| C<sub>max</sub> | Geom. Mean [hng/mL] | 2566.481 | 1234.14 | 1373.78 | 74.62 | 19.38 | 8.07 |
| t<sub>max</sub> | Median (range) [h] | 4 (1-6) | 1 (1-6) | 4 (1-8) | 2 (2-3) | 2 (1-2) | 2 (2-3) |
| t<sub>lag</sub> | Median (range) [h] | 0.5 (0.5-3) | 0.5 (0.5-4) | 0.5 (0.5-4) | 0.5 (0.5-0.5) | 0.5 (0.5-0.5) | 0.5 (0.5-0.5) |
| t<sub>1/2</sub> | Mean (SD) [h] | 3.09 (1.11) | - | 1.97 (0.26) | 1.30 (0.23) | 2.05 (1.574) | 2.05 (0.306) |
| Ratio fed /<br>fasted | AUC<sub>(0-last)</sub> | 1.12 | 23.36 | 0.90 | 1.33 | 0.69 | 1.04 |
|  | C<sub>max</sub> | 0.92 | 5.77 | 0.92 | 1.13 | 0.48 | 0.75 |

**Figure 1:**
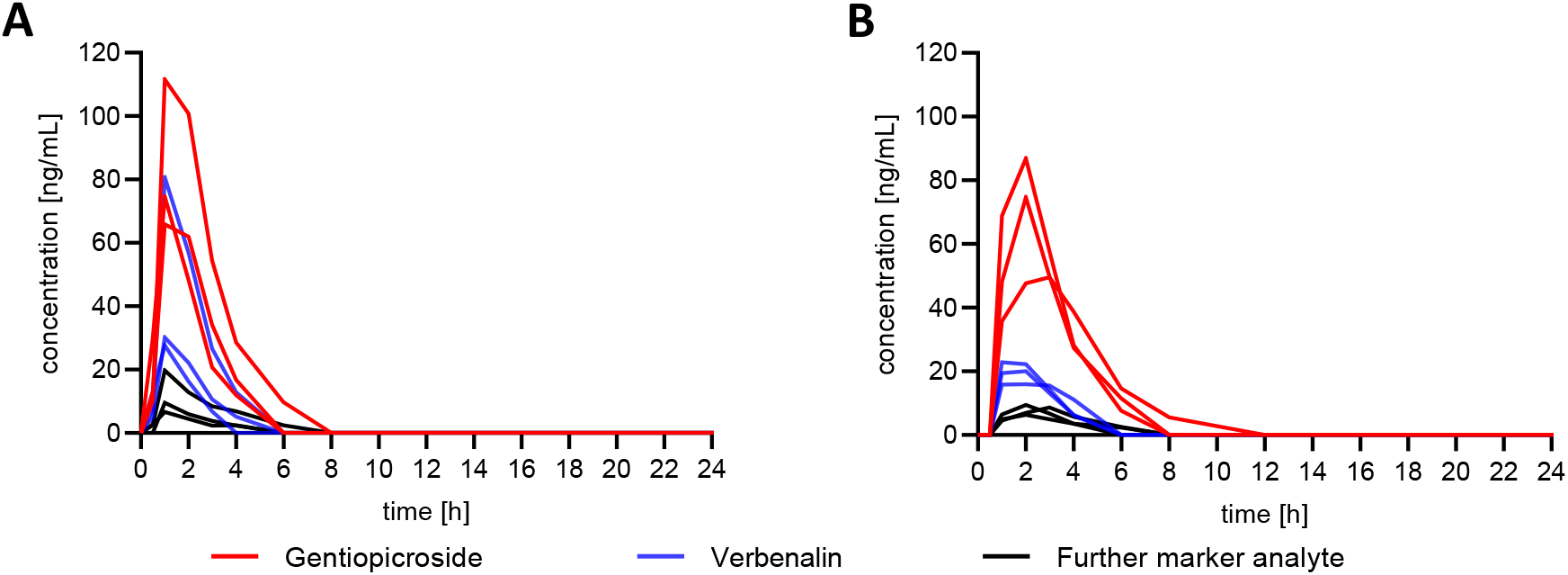
Overlay of individual plasma concentration *vs.* time curves of gentiopicroside, verbenalin and a further marker analyte in dogs (N=3) after administration of BNO 1016 under fasting (A) and fed (B) conditions.

**Figure 2:**
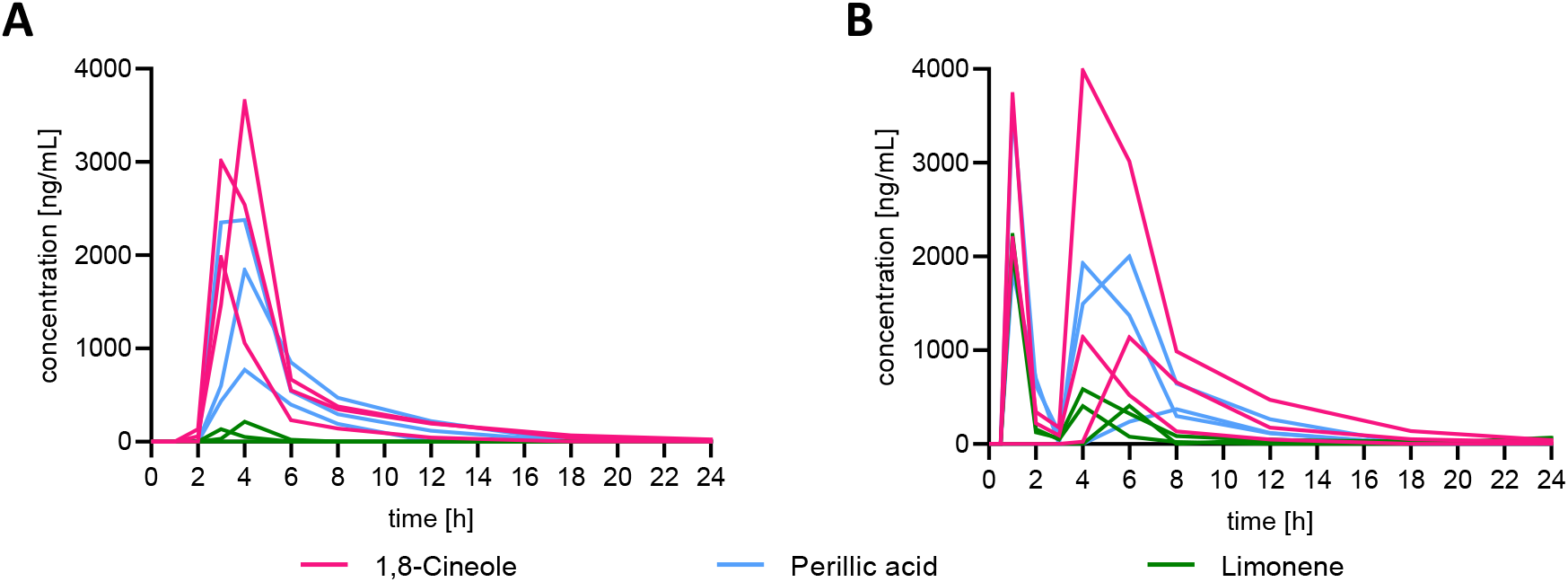
Overlay of individual plasma concentration vs. time curves of 1,8-cineole, perillic acid and limonene in dogs (N=3) after administration of ELOM-080 under fasting (A) and fed (B) conditions.

**Figure 3:**
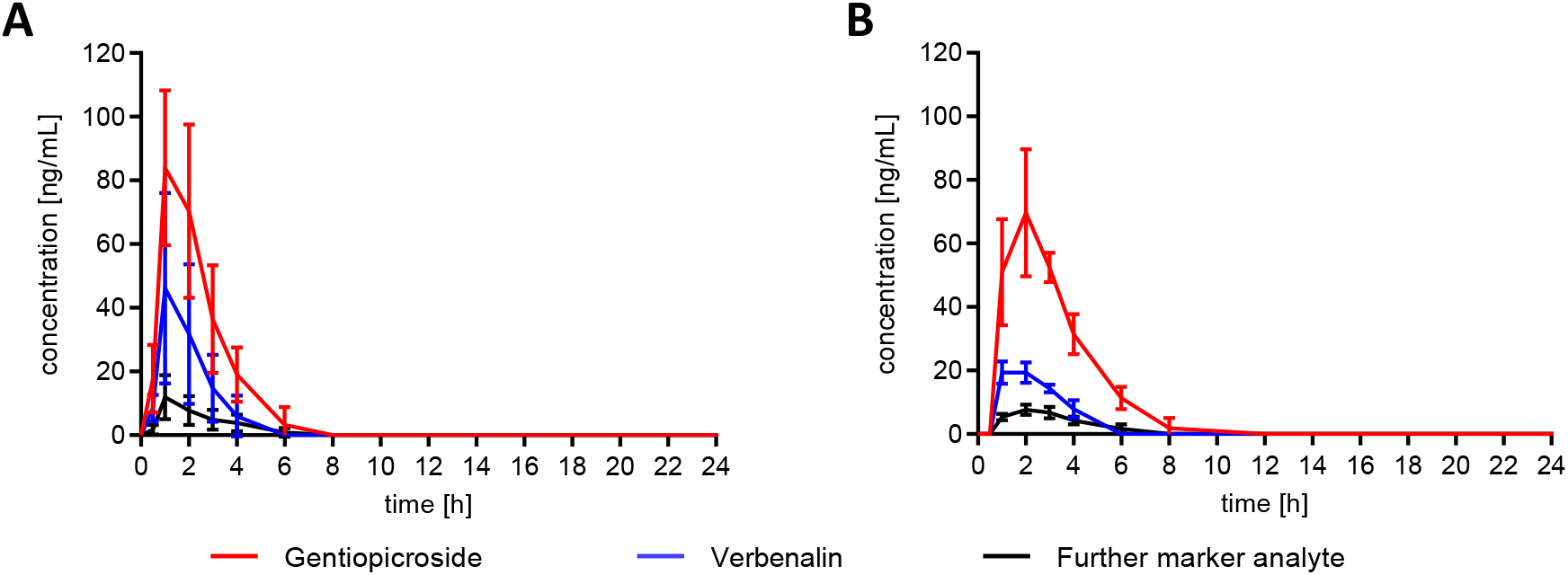
Mean plasma concentration vs. time curves (±SD) of gentiopicroside, verbenalin and a further marker analyte in dogs (N=3) after administration of BNO 1016 under fasting (A) and fed (B) conditions.

**Figure 4:**
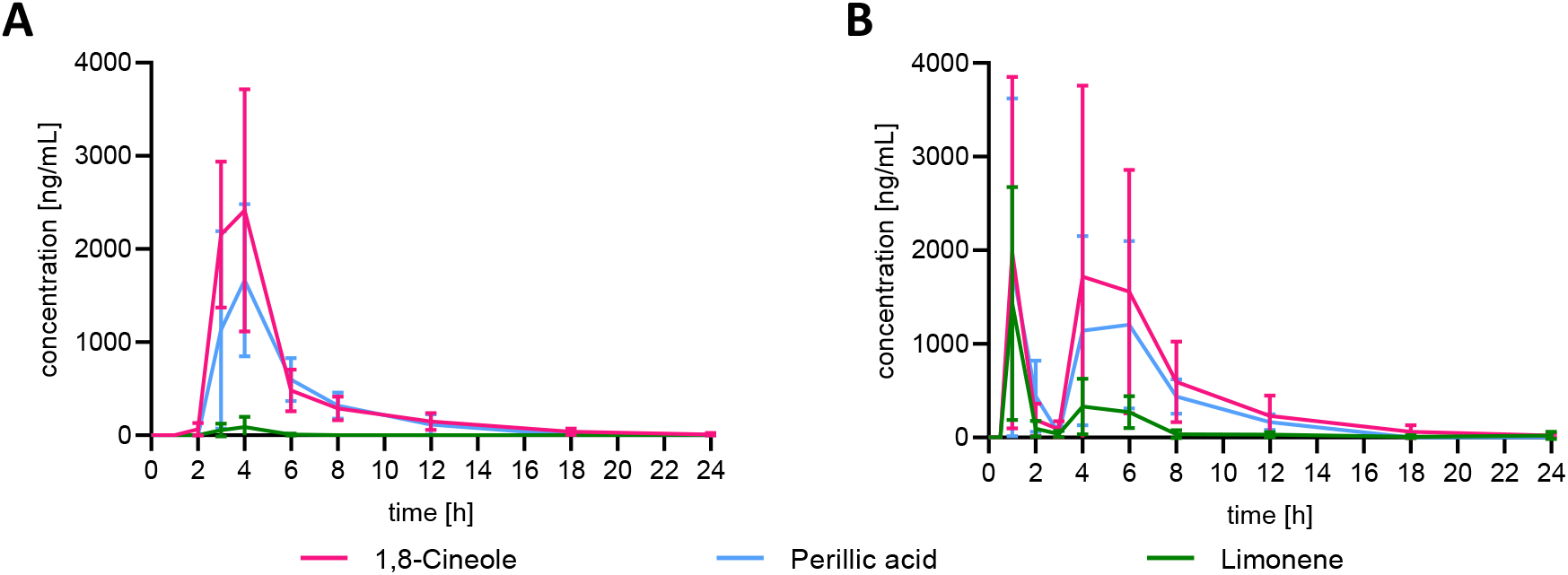
Mean plasma concentration vs. time curves (±SD) of 1,8-cineole, perillic acid and limonene in dogs (N=3) after administration of ELOM-080 under fasting (A) and fed (B) conditions.

In general, the three analytes determined for BNO 1016 demonstrated a homogenous course in the fasted as well as in the fed state. Typical characteristics of an immediate-release drug formulation with a rapid increase until C_max_ with median t_max_ values of 1 h in the fasted and 2 h in the fed state could be observed for all three analytes detected. Curves showed the typical sigmoidal course after single dose administration (Figure 1). There was no lag-time (t_lag_) under fasting and only an expected slight delay of 0.5 h under fed conditions. The rate and extent of exposure (C_max_, AUC_(0-tlast)_) of verbenalin decreased after intake of food (ratio of geometric means fed/fasted: 0.48 and 0.69), whereas that of the further marker analyte was nearly unchanged (ratio fed/fasted: 0.75 and 1.04). Geometric mean ratios of C_max_ and AUC_(0-tlast)_ values of gentiopicroside were slightly increased with 1.13 and 1.33, respectively. Mean apparent elimination half-life times (t_1/2_) of the three analytes lay in the range of 0.96-1.78 h (fasted) and 1.30-2.05 h (fed), suggesting that elimination kinetics are unchanged (see Table 1). Overall, a relevant food effect could not be observed after administration in dogs.

Individual overlays of the ELOM-080 analytes in the fasted state also showed an overall homogeneous picture in the three dogs (Figure 2). Interestingly, lag-times of up to 2 h post administration could be observed for at least two of the three analytes, reflecting a longer residence time of the formulations in the dogs’ stomach until drug liberation in the intestine. Corresponding t_max_ values lay in the range of 3 to 4 h *p.a.*. The individual overlays after the administration of the high fat diet show two major peaks. This occurred in two of the three animals. A sigmoidal course after administration can at best be guessed for the second major peak of the individual curves. Observed median t_lag_ was 0.5 h but ranged from 0.5 to 4 h for all analytes. Raw data of one of the three dogs revealed only 1 peak for all analytes with tmax values up to 8 h and t_lag_ of 4 h. C_max_ and AUC_(0-tlast)_ of 1,8-cineole (ratio of geometric means fed/fasted: 0.92 and 1.12) and perillic acid (ratio of geometric means fed/fasted: 0.92 and 0.90) were in the same magnitude under fasted and fed administration, whereas that of limonene was increased (ratio fed/fasted: 5.77 and 23.36). Mean apparent elimination half-life times of the analytes lay in the range from 2.98 to 3.51 h (fasted) and 1.97 to3.09 h (fed). Although t_1/2_ of limonene could not be derived properly, the data suggest that elimination kinetics are not altered by food intake, as expected (see Table 1).

## Discussion

The present study investigated the pharmacokinetics of BNO 1016 and ELOM-080 with special attention to (1) the rate of absorption and (2) the occurrence of a food effect depending on preceding feed intake in male beagle dogs.

Both herbal medicinal products are widely used in Germany and Europe for the treatment of acute rhinosinusitis. BNO 1016 is available as an immediate release formulation (coated tablet) and ELOM-080 is contained in an enteric-coated soft gelatin capsule. It can be assumed that absorption of active pharmaceutical ingredients (APIs) from BNO 1016 occurs fast and therefore, results also in a fast onset of treatment effects. The purpose of an enteric coating is to delay dissolution and the release of APIs from the drug formulation until it has reached the intestine. Therefore, it might be suggested that retention time of an enteric coated soft gelatin capsule in the stomach is influenced when food has been consumed prior to ingestion or immediately afterwards. The time indigestible solid particles are retained in the stomach usually depends on the diameter of the dosage form, the frequency of food intake, as well as on the composition and caloric density of an administered meal [20][21]. In humans, published data show that single unit enteric-coated dosage forms may remain in the stomach for 10 h or even longer when co-administered with a high fat breakfast followed by several meals over the course of the day [22].

Moreover, it is known that gastric emptying of drugs may be even delayed for up to 30 h in non-fasted animals [23].

In this first pilot study, we chose beagle dogs as the standard model to study food effects [19]. Based on our previous experience with the analytical detectability of analytes in plasma from rats and dogs, we chose to administer the 5-fold equivalent of the recommended human daily dose assuming that the physiological performance of the single formulations will not change with the dosage administered. Plasma samples were withdrawn until 30 h *p.a.* to avoid missing possible late-occurring absorption processes. In general, the analytes determined for BNO 1016 demonstrated a homogenous course of the individual plasma concentrations *vs*. time curves in the fasted as well as in the fed state. Lag-times of the absorption process lay in the expected range of 0-0.5 h *p.a.* for an immediate release formulation with corresponding median t_max_ values of 1-2 h. A relevant food effect could be ruled out. For the enteric-coated formulation of ELOM-080 however, we observed lag-times of the absorption up to 2 h *p.a.* even in the fasted state, reflecting longer gastric residence times of the formulations in the dogs. In the fed state a pronounced food effect could be noticed with median lag-times up to 3-4 h *p.a.* and a splitting into two major peaks in plasma curves in two of the three dogs. The cause of this double peak phenomenon is unknown. Neither the inspection of the laboratory journals nor the direct questioning of the staff gave any indications of misadministration, *e.g.* dogs biting capsules etc.. However, the first peak could be a sign for a bursting of soft gelatin capsules in the stomach with direct release of the active ingredients.

While, according to the respective SmPC [4], the intake of ELOM-080 is recommended before meals, a truly empty stomach in humans can usually only be expected in the mornings after overnight fasting. The consumption of meals and snacks throughout the day might practically lead to a closed pylorus sphincter over long time distances during the day [24]. A respective food effect in humans for ELOM-080 should not lead to severe safety-relevant side effects. However, if this finding in dogs translates to humans, some of the frequently observed gastrointestinal side effects, like stomach and upper abdominal complaints [4], might be explained. Most certainly, a precise dosing with a rapid release and onset of action does not seem likely based on the data obtained in the present study. Therefore, it seems quite valuable to verify these findings in a further pharmacokinetic study in humans, considering that patients suffering from respiratory tract disorders take ELOM-080 with the expectation of a fast relief of their complaints.

*In conclusion*, our pilot study provides evidence that, in contrast to the immediate release medicine BNO 1016 with fast and homogenous absorption characteristics, absorption of ELOM-080 from an enteric-coated capsule might be substantially affected by food intake in humans. Consequently, a faster and more precise onset of action in humans can be assumed for BNO 1016.

## Abbreviations

API: active pharmaceutical ingredient
AUC_(0-tlast)_: area under the plasma concentration *vs*. time curve from dosing time to the last measurement time point with a concentration value above the lower limit of quantitation, calculated by means of the linear/log trapezoidal method which uses the linear trapezoidal rule up to C_max_ and afterwards the log(interpolation) trapezoidal rule for the subsequent part of the curve
C_max_: maximum concentration in plasma, directly taken from measured concentration values
EMA: European Medicines Agency
GC-MS: gas chromatography coupled to mass spectrometry
LC-MS/MS: liquid chromatography coupled to mass spectrometry
LLOQ: lower limit of quantification
MMC: migrating motor complex
*p.a.*: post administration
SmPC: Summary of product characteristics
t_lag_: time from administration to first quantifiable time point in plasma
t_max_: time to reach maximum concentration, obtained directly from measured values
t_1/2_: apparent terminal elimination half-life

## Acknowledgements

We thank Dr. Damaris Kukuk, at Eurofins BSL, Munich, Germany and Lucile Gioda at Avogadro LS, Fontenilles, France for project management and conduct of the in-life phase of the animal study as well as Dr. Bernhard Nausch for critical reading of the manuscript.

## Author’s contributions

Jan Seibel made substantial contributions to conception and interpretation of data and has been involved in drafting the manuscript. Meinolf Wonnemann has made substantial contributions to the conception and interpretation of data and has been involved in pharmacokinetic calculations as well as drafting the manuscript. Astrid Neumann and Anne Müller have been involved in the bioanalytical planning, performance and bioanalytical evaluation.

